# Affinity selection in germinal centers: Cautionary tales and new opportunities

**DOI:** 10.1101/2021.03.12.435085

**Authors:** Jose Faro, Mario Castro

## Abstract

Our current quantitative knowledge of the kinetics of antibody-mediated immunity is partly based on idealized experiments throughout the last decades. However, new experimental techniques often render contradictory quantitative outcomes that shake previously uncontroversial assumptions. This has been the case in the field of T-cell receptors, where recent techniques for measuring the 2-dimensional rate constants of T-cell receptor-ligand interactions exposed contradictory results to those obtained with techniques measuring 3-dimensional interactions. Recently, we have developed a mathematical framework to rationalize those discrepancies focusing on the proper fine-grained description of the underlying kinetic steps involved in the immune synapse. In this perspective article, we apply this approach to unveil potential blind spots in the case of B-cell receptors (BCR) and to rethink the interactions between B cells and follicular dendritic cells (FDC) during the germinal center (GC) reaction. Also, we elaborate on the concept of “catch bonds” and on the recent observations that B-cell synapses retract and pull antigen generating a “retracting force”, and propose some testable predictions that can lead to future research.

## 1. Introduction

The history of Immunology runs in parallel to the use of innovative experimental techniques. From the early microscopes to modern molecular-level sensors, our basic knowledge has refined through the years; new techniques provide new discoveries. For instance, in 2010 two papers [1,2] revealed essential differences between traditional assays for the interaction of T-cell receptors (TCR) and their peptide-major histocompatibility complex (pMHC) ligands, in which one of the reactants is in fluid phase (referred to as 3-dimensional (3D) assays), and those in which both TCRs and pMHC ligands are confined to a cell membrane (referred to as 2D assays). More recently, an additional layer of complexity was added to the binding kinetics of TCRs with the discovery that some TCRs display a catch-slip bond behavior when are bound to a given pMHC ligand [3,4]. While the predominant binding of receptors to ligands, known as slip bonds, are characterized by being weakened when a tensile force is applied between the receptor and the ligand, such that the half-life of the bond decreases for increasing tensile force [5,6], in catch bonds the half-life increases for an increase of the tensile force, and catch-slip bonds display initially a catch bond behavior until a threshold value is reached, beyond which the bond becomes a slip one and its half-life decreases for increasing force [6,7]. Typically, the optimum tensile force for TCR-pMHC interactions with catch bond behavior is in the range of 10-20 pN [4], which is within the range of forces per bound TCR generated by pulling in cell-cell interactions [8] and during initial TCR-pMHC binding [9]. Currently, although there is no consensus about the mechanical basis of the catch bond behavior of TCR-pMHC interactions, there is consensus about the role of dimensionality and on how different experimental methods can dramatically change previous qualitative and quantitative predictions. In principle, this is how normal science progresses, but when a dominant conceptual paradigm is built based on inappropriate experimental methods it prevents moving the field forward (an illustrative example is the case of the oligoclonal seeding concept in germinal center studies [10,11]). In these cases, mathematical modeling can contribute to unveil important differences between experimental procedures, to warn against intuitively appealing but wrong interpretations of experimental results, and also to suggest new venues to explore [12].

Recently, we have developed a theoretical framework to analyze the effective binding rate constants of TCR-pMHC interactions in 3D and 2D assays [13]. The rationale behind our approach is to disentangle kinetic mechanisms into fine-grained steps. For instance, in that framework binding-unbinding is understood as a sequence of three independent mechanisms: diffusion (translation), orientation (rotation), and molecular binding. Our approach successfully helped to reconcile contradictory experimental results as well as to predict how different experimental assays would lead to divergent predictions [13].

In the case of antibodies (Ab) and B-cell membrane immunoglobulins (Ig), here denoted also as B-cell receptors (BCR), there are also potential pitfalls in interpreting experimental data. As already acknowledged years ago by Mason and Williams [14] the interaction of bivalent Abs to antigen in solution departs considerably from their interaction with cell-bound antigen. Moreover, it has been soundly argued that in real ligand-receptor reactions it is essential to understand the process of events in dynamic cell-cell interactions rather than following equilibrium considerations [15]. This raises the question of how many of the current underlying assumptions on the binding of BCRs to an antigen (Ag) displayed on cell membranes are based on misinterpreted experimental results or on inappropriate extrapolation from in vitro fluid phase systems to very different conditions such as cellular systems.

In this perspective article, we apply our theoretical framework to illustrate how it can help to rightly understand and compare binding results of Ab-Ag reactions in solution (3D assays) and in cell-cell interactions during an immune respoonse (2D interactions, as it can be expected *in vivo*). Specifically, we will approach the following two questions: (i) does the dimensionality of the experimental setup impinge in our understanding of interactions of BCRs and Ags held on the membrane of follicular dendritic cells (FDC) during a germinal center (GC) reaction?; (ii) is there a role for catch-bonds in BCR-Ag reactions, and how it could affect Ab affinity maturation in GC reactions?

## 2. General theoretical framework and derived properties for antibody- and BCR-ligand interactions

Binding and unbinding of ligands (*L*) to receptors (*R*) can be described in detail as a series of independent steps. The first step comprises the spatial approximation of *R* and *L* molecules to a distance that allows binding (denoted by *RL*\*). Once in that *state*, the two molecules must attain a proper orientation to bind to each other —we denote this intermediate-oriented state as *RL*— until finally, they can form a complex *C* [13]. We summarize these series of states and processes in Eq. (1), where the translational and rotational diffusion on and *off* constant rates correspond either to those in 3D or in 2D settings.

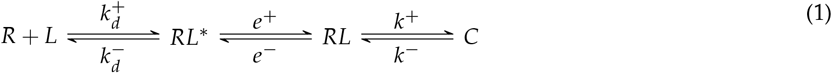

### 2.1. *Experimentally estimated effective* on *and* off *constant rates*

As we have shown previously [13], in the cases of the Surface Plasmon Resonance (SPR, a 3D assay) and the Adhesion Frequency assay (a 2D assay [1]) Eq. (1) is effectively simplified to a single-step description of the process, namely,

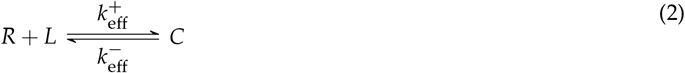

In these two cases the effective binding and unbinding rate constants, respectively, 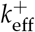 and 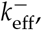 are given by the following expressions [13]:

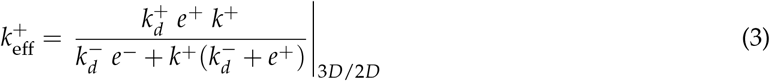

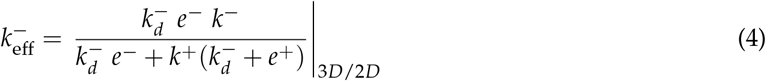

where the subscripts indicate that the rate constants are those corresponding to either 3D or 2D reactions, and *k*^+^ and *k*^−^ are the truly *on* and *off* constant rates, which are, therefore, identical for the 3D and the 2D interactions.

The effective association affinity constant is simply the product of the affinities (denoted by *capital K*) of every intermediary step:

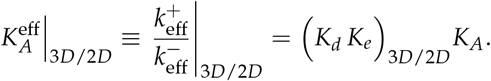

When this formalism is applied to Ab-Ag and BCR-Ag interactions it is instructive to plot 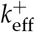 as a function of *k*^+^, and 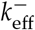 as a function of *k*^+^ and *k*^−^ because it shows how misleading can be the estimation of *k*^+^ and *k*^−^ by the effective rate constants 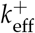 and 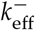, respectively. For instance, according to Eq. (3), for Abs and BCRs of the same isotype, when 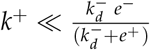 the effective rate 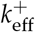 will increase linearly for increasing values of *k*^+^. However, when *k*^+^ approaches the value 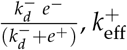 will increase more and more slowly, approaching asymptotically the maximum 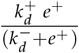 when 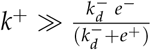 (see Fig. 1A). The latter situation could be named diffusion/orientation-limited reactions as the limiting process is not binding itself but getting the Ab or BCR and the Ag close enough and well oriented before the real molecular binding occurs.

**Figure 1.**
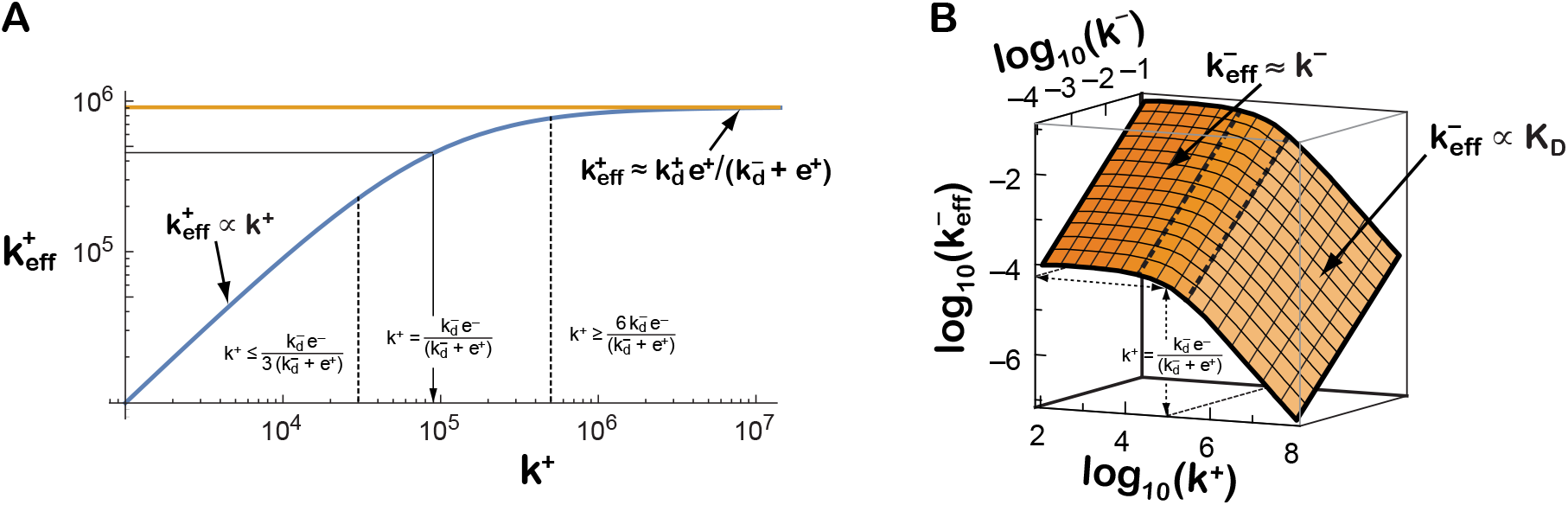
Effective kinetic rates measured experimentally can be misleadingly interpreted as approximations of the underlying molecular binding constant rates. Expected dependency of 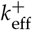 (panel **A**) and 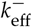 (panel **B**) as a function of *k*^+^ and *k*^−^, for a given set of values of the translational and rotational rate constants 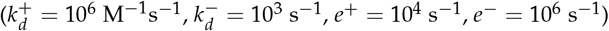.

With respect to 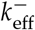, according to Eq. (4), when 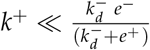 for any given value of *k*^+^ the effective off rate constant is a good estimation of *k*^−^. However, for values of 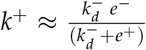 this estimation is not longer valid, and for 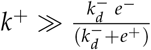 the value of 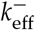 becomes instead a good estimation of the true dissociation constant, more precisely 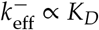 (see Fig. 1B). In the remaining of this paper we refer to the quantity 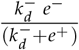 as the *k*^+^-threshold.

In summary, the following approximations hold when *k*^+^ is greater or smaller than 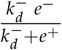:

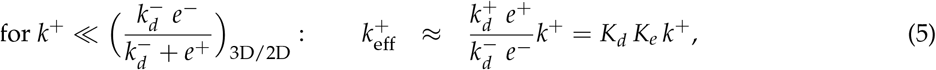

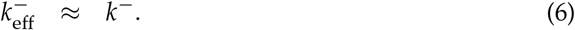

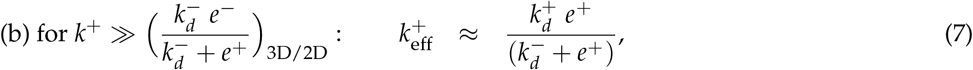

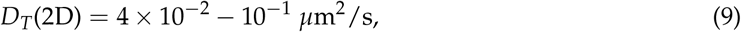

In the following sections we discuss succinctly the mechanisms and geometrical aspects that might raise concerns about our understanding of Ab-Ag and BCR-Ag interactions: diffusion, rotation and dimensionality.

### 2.2. The role of translational diffusion

Eqs. (3)–(4) and (5)–(8) are the roadmap to unveil the differences imposed by geometrical, physical and molecular constraints. In particular, due to geometrical constraints, translational diffusion *on* and *off* constant rates are mathematically different in 2D (membrane) and 3D (solution). Specifically, for unbound BCRs of IgM and IgG isotypes the median translational diffusion constant has been estimated to be [16–18]

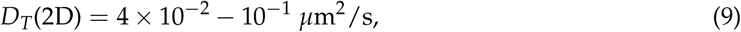

and for IgD BCRs *D_T_*(2D) = 3 × 10 ^−3^ *μ*m^2^/s [17]. However, for free IgMs and IgGs in diluted saline solutions, as is the case in many SPR experimental setups, *D_T_*(3D) has been reported to be [19–21]

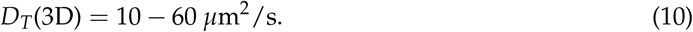

In a 2D setting, the translational diffusion *on* rate constant of BCRs toward an Ag tethered on a cell membrane can be approximated by the expression: 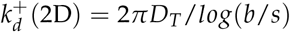, where *D_T_* is the translational diffusion constant of BCRs in the membrane, *b* is the the mean distance between BCRs in the membrane of a B cell, and *s* is the encounter radius, which is essentially the radius of a BCR (~7.5 nm [22]). Following Lauffenburger and Linderman (see p. 153 in Ref. [23]), to convert 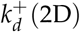 into the same unit as a typical bimolecular *on* rate, we multiply by *N_A_* and, to give an estimated local volume for cell surface components, by a membrane thickness of *ℓ* = 10 nm. Considering a B cell with a surface area of 40 *μ*m^2^ (corresponding to a diameter of ~ 7*μ*m) and about 50,000 BCRs, it follows b ≈ 28 nm. Then, using *D_T_* from Eq. (9) one finds:

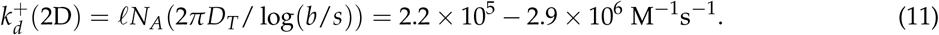

The corresponding translational diffusion *off* rate constant can be approximated as:

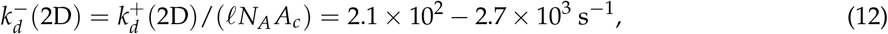

where *A_c_* is the effective local cell surface area, defined as *A_c_* = *πs*^2^ [23].

In 3D the translational diffusion *on* and *off* rate constants of proteins can be approximated by 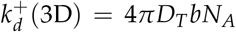 and 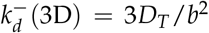 [13]. This applied to free IgGs using the estimate in Eq. (10) yields:

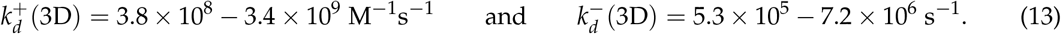

### 2.3. The role of rotational diffusion

Antibodies and BCRs bind to Ags through a small part in the variable region, a small area at the end of each Fab arm known as antigen-binding site or paratope (*P*). The corresponding region at the surface of the cognate Ag, that is, the actual part of the Ag participating in the Ab-Ag or BCR-Ag bond, is denoted epitope (*E*).

The spatial position of a BCR Fab paratope is determined by two possible Fab twists and two possible Fab tilts, sketched in Figs. 2a and 2b. The twists are: 1) of a whole BCR relative to the axis along the Fc region, normal to the plasma membrane, denoted here Fc twist rotation, with values in principle in the range [-360°, +360°]; and 2) of a Fab arm relative to its major axis (i.e., the hinge-paratope axis), denoted here Fab twist rotation, with values, at least for some Ig isotypes, in the range [−180°, +180°] relative to the more relaxed conformation [24,25].

**Figure 2.**
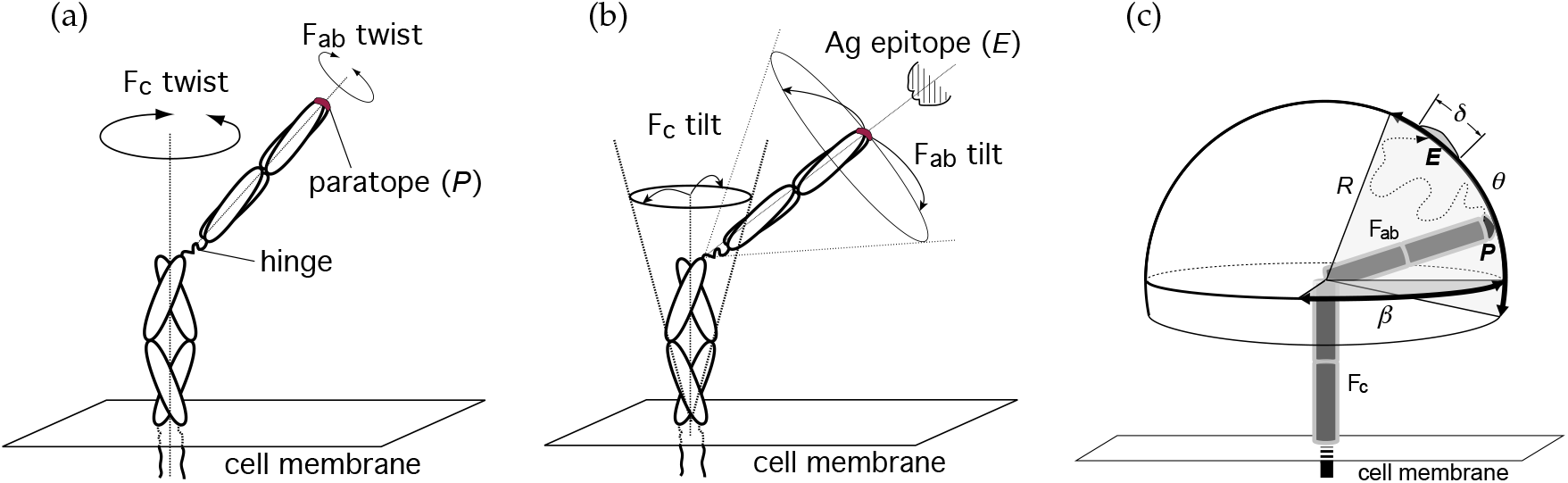
Possible rotations having an impact on the position of a paratope *P* relative to an Ag epitope *E*. The position of *P* in a membrane Ig is submitted to (a) two possible twists, and (b) two possible tilts (wagging and wobbling). In (c) it is shown a sketch of the diffusion of *P* constrained to a spherical cap on a sphere of radius *R* = length of a Fab arm ≈ 84 Å [29] and to Fab altitude and azimuth angles *θ* and *β*, respectively. Depicted is a possible path followed by paratope *P* toward epitope *E* (dotted line). This movement corresponds to Fab tilt in (b). The angle *δ* corresponds to an arc on the spherical surface equal to the major length of *E*.

Similarly, the tilts or wagging and wobbling are: 1) of the whole BCR relative to the major axis of the molecule (along the Fc region, normal to the plasma membrane) denoted here Fc tilt; and 2) of a Fab arm relative to its major axis, denoted here Fab tilt; the range of this tilt is much larger than the Fc tilt range (in some isotypes the angle of the Fab tilt cone may be even larger than 90° [26–28]).

Considering the rotational movements sketched in Fig. 2 a and b, once a BCR and an Ag are in the *RL*^*^ state one can map the binding problem to that of random walk of a paratope on a region of a spherical surface of radius *R* = *length of a Fab arm* ≈ 84 Å [22,29] delimited by altitude and azimuthal angles *θ* and *β*, respectively, constrained by the flexibility of the hinge of the Fab arm (Fig. 2c). Thus, a paratope *P* diffuses within this restricted spherical surface region until it reaches its cognate epitope *E*. This epitope can be described as a small area on the mentioned spherical surface covering an arc of maximum length *L* ≈ 28 Å [30], which is equivalent to an angle *δ* ≈ 19° (Fig. 2c). In addition, once the paratope *P* is close to the epitope *E*, proper binding requires their proper alignment or orientation through Fab twist rotation, so not only rotational-tilt diffusion to the right location matters, also proper alignment is mandatory (twist or rotational-alignment). Note that following unbinding the possibility of immediate rebinding is lost when either movement is reversed, namely, the alignment is lost or the Fab paratope *P* leaves the proximity of the epitope *E* through tilting diffusion.

The different IgG subclasses in human and mouse have different hinge regions, and this makes them differ substantially in the flexibility and rotation of their Fab arms with respect to the Fc region (tilt rotation) [31,32]. More specifically, it has been found that human IgG subclasses can be ordered with respect to hinge flexibility between Fab arms (range of Fab-Fab angles), from most to least flexible, as: IgG3 > IgG1 > IgG4 > IgG2 [26]. Also, it has been shown that murine IgG subclasses can be ranked with respect to their Fab arms flexibility as follows: IgG2b > IgG2a > IgG1 > IgG3 [33]. This two orderings turn out to be totally consistent with each other when the reported functional equivalence of murine and human IgG subclasses is considered [34,35]. For instance, murine IgG2b corresponds to human IgG3 [34] and the maximum tilting angle (Fab-Fc angle) reported in the literature is about *θ* ≃ 70° for both [26,33]. Similarly, murine IgG1 corresponds to human IgG4 and both have *θ* ≃ 50°. These differences and their impact on the average speed of the tilt Fab rotation [33], suggest that Ig class switching could alter the effective binding constant rates of BCR-Ag and Ab-Ag interactions because of a concomitant change in their paratope rotational diffusion rate constants.

### 2.4. The role of dimension

Using experimental data on relaxation times of fluorescent polarization anisotropy of fluorescent probes [33,36] quantitative information on molecular rotational velocity can be obtained. For instance, the mean square angular deviation of a molecule rotating about a given molecule’s axis after a time *t* is given by [37,38]

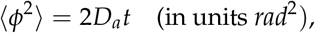

where *D_a_* is the corresponding twist (alignment) rotational diffusion constant. On the other hand, the mean rotational correlation time of a molecule rotating about a given molecule’s axis, here denoted *t*_1rad_, is the time it takes on average to rotate 1 rad. From these two definitions it follows that for a Fab arm rotating about its major axis (Fab twist) the twist (alignment) diffusion constant is:

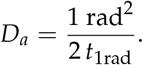

However, for a Fab arm rotating about the Fc axis with center in the hinge region (Fab tilt) the tilt rotational diffusion constant is [36]:

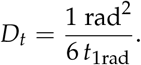

The values of *t*_1rad_ corresponding to Fab tilt as well as to Fab twist have been determined for IgG molecules from fluorescent anisotropy experimental data, with IgG molecules both in solution (*t*_1rad_(3D)) and attached through the Fc to a large particle (*t*_lrad_(2D)) [33,36,39,40]. Based on that data it can be estimated that the corresponding 3D rotational (tilt+twist) *off* constant rates *e*^−^ are at least 3 orders of magnitude faster than those for 3D translational diffusion *off* constant rates 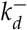 (Eq. (13)). However, in 2D the *e*^−^ constant rates are even much faster than the 2D 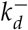 constant rates (Eq. (12)). Thus, the *k*^+^-threshold values in 3D and in 2D can differ from each other by several orders of magnitude. This points out, again, to the need of carefully estimate these rates in order to properly understand the outcome of different experimental setups.

In general, for Ab-Ag interactions in a typical experiment using SPR (a 3D assay), the translational constant rates and the rotational constant rates will depend on which of the two interacting molecules is immobilized. There are two possibilities: (1) antigens in solution and Abs immobilized, (2) antigens immobilized and Abs in solution. In the first case, the translational constant rates, 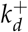 and 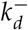, will be only slightly different for Ags of widely different sizes and shapes [20], but the rotational diffusion constants *D_t_* and *D_a_* will be very different for Ags of widely different sizes and shapes, and different (but not very different) for Abs of different isotype or subclass. As a consequence, the total rotational diffusion constants (*D_t_* of Ag + *D_t_* of Ab, and *D_a_* of Ag + *D_a_* of Ab) can be very different for different Ags and Abs, and therefore can differ very much in their rotational rate constants *e*^+^ and *e*^−^. In contrast, in the second case (Ag immobilized, Ab in solution), the translational diffusion constant and the corresponding rate constants, 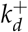 and 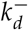, will be very similar for Abs of different isotypes [20], and the rotational rate constants, *e*^+^ and *e*^−^, will be the same for Abs of same isotype and subclass, and different (but not very different) for Abs of different isotype or subclass.

Therefore, according to Eqs. (3) and (4), the differential contribution of Abs and Ags to the 3D rotational constant rates of Ab-Ag interactions is expected to have an important impact on the effective 3D rate constants, 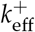 and 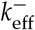 depending on which of the two interacting molecules, Abs or Ags, is immobilized, a theoretical prediction consistent with experimental observations [41]. However, in 2D reactions BCRs interact with Ag in form of immune complexes tethered on cell membranes, notably FDCs. These Ag complexes (termed iccosomes [42]) are much larger than BCRs (25 to 50-fold difference in size) and are anchored to the FDC membrane by several receptor molecules, namely, Fc receptors and C3b/C4b receptors [22,42] (Fig. 3). Hence, the contribution of Ag complexes to diffusion and rotation are expected to be negligible compared to that of BCRs. Then, for different BCRs and Ags the translational rate constants, 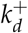 and 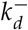, would be similar because BCRs of different isotypes have similar 2D translational diffusion constants [16–18], and the rotational rate constants, *e*^+^ and *e*^−^, would be similar for BCRs of same isotype/subclass, and different (but not very different) for BCRs of different isotype or subclass.

**Figure 3.**
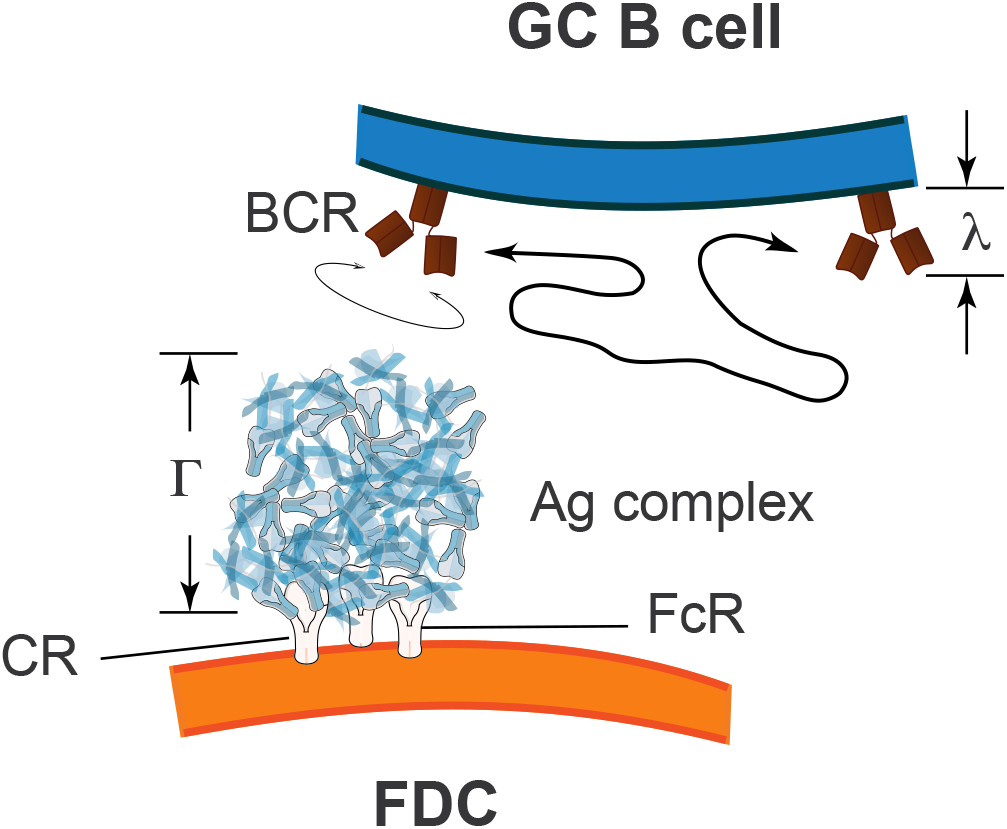
BCRs contribute to translational and rotational diffusion to a much larger extent than Ag complexes on FDCs (termed iccosomes). Iccosomes are tethered to the FDC membrane through several Fc and/or complement receptors. Typical diameter of iccosomes is Γ = 250-700 nm [42], while BCR diameter is *λ =* 10-15 nm [22]. FcR, Fc receptor; CR, complement receptor.

In general, all the above indicates that for Ab-Ag interactions the *k*^+^-threshold value 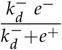 in 3D assays can vary substantially, particularly for Ags of different size when they are in solution. However, the corresponding *k*^+^-threshold value in 2D BCR-Ag interactions will be independent of Ag (save for epitope valency) and of BCRs, except for BCRs of different isotype and subclass, which can have *k*^+^-threshold values differing by at most one order of magnitude.

## 3. Is there a role for catch-bonds in BCR-ligand interactions?

### 3.1. Evidence for mechanical forces in B-cell synapses and their relationship to kinetic rate constants

The binding of BCRs to Ag tethered to a plasma membrane induces in B cells the formation of an immune synapse. After the synapse is formed, an initial lateral traction force acts on Ag-bound BCRs, followed by a later retraction that pulls the Ag-bound BCRs [43–45]. The formation of such immune synapses involves an initial clustering of Ag-bound BCRs by an active process, driven intracellularly by myosin, F-actin and dynein [44,46], that generates shear or lateral forces of 10-20 nN per 50 *μ*m^2^ acting on Ag-bound BCRs [46]. Assuming that in that large area there are initially about 10^3^ Ag-engaged BCRs, individual BCR-Ag bonds would be submitted to a shear force of 10-20 pN that would increase as the number of broken BCR-Ag bonds increase. The subsequent immune synapse retraction pulls the Ag-bound BCRs, generating a tensile force of 10-20 pN per Ag-bound BCR in the direction normal to the local membrane plane [18,44].

For the process of Ag pulling and extraction, of the three processes and four states distinguished above in BCR-Ag interactions only unbinding and the bound state are relevant. For instance, in order for GC B cells to be able to pull and extract Ag from FDC membrane immune complexes the average time 1/*k*^−^ has to be large enough compared with the time scale required for the Ag “pulling and extracting” process. This time scale is similar to the time spent by a B cell interacting with a FDC, which has been reported to be on average less than 5 min [47]. Assuming this time is 5 min, 1/*k*^−^ has to be larger than 300 s, or *k*^−^ < 3 × 10^−3^ s^−1^, to allow for Ag extraction. Nevertheless, for larger values of *k*^−^, if compensated by very large values of 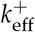, it could still be possible for a B cell to extract Ag.

### 3.2. An extension of the theoretical framework

It has been shown that some cell-receptor adhesion molecules can be in two different conformational states, the so-called bent and extended states, the second one being favored in receptor-ligand complexes submitted to a tensile force [48]. Also, recently it has been described that TCRs can display dynamic ectodomain allostery [8,49]. Analogously, it has been recently reported that BCRs binding to Ag tethered on a plasma membrane undergo allosteric changes in the CH2 domain, in the B cell membrane-proximal CH domain, as well as in the cytoplasmic tail of IgG BCRs [16,18,50]. Similarly, Ag binding by IgG Abs induced a conformational change in the Fc region of the Abs [51]. Furthermore, a recent analysis of the structure of 15 anti-protein Abs comparing the antigen-bound *vs* unbound forms has revealed that the binding process induced in a large fraction of those Abs high allosteric changes in the angle between the variable and the adjacent constant domains in both the heavy and light chains [52]. Finally, it has also been shown using different approaches that the paratope of Abs and BCRs can be at least in two different conformational states [53,54].

Considering such conformational flexibility of the paratope of BCRs and Abs and the allosteric changes often brought about by antigen binding we assume that in general the complex *C* in Eq. (1) comprise in fact at least two different complex states, *C*_0_ and *C*_1_, which can be affected allosterically or in some other form by an applied force. Without loss of generality, we assume that in the absence of any applied force the state *C*_0_ is more probable than the state *C*_1_. Eq. (14) summarizes all the states and reactions for most 3D experimental settings in which a free ligand (or receptor) diffuses until it binds to a surface linked receptor (or ligand).

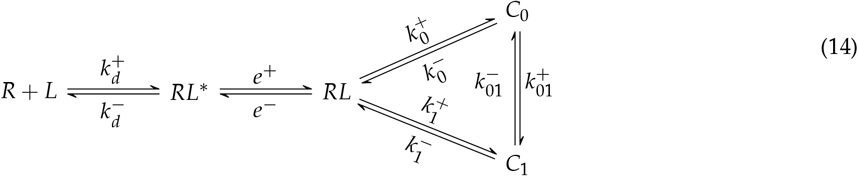

with 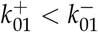.

If Ab class switch induces a dominant conformation swap between the *C*_0_ and *C*_1_ conformations, this can be expressed in terms of Eq. (14) as a change in the constant rates 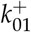 and 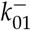 to a new ones, 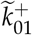 and 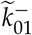, respectively, such that 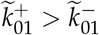. This possibility could have important consequences when considering the effect of lateral and pulling forces on BCR-Ag bonds.

The situation in B cell-FDC interactions and in some experimental 2D settings is slightly different from that shown in Eq. (14). For instance, in a variant of the Biomembrane Force Probe (BFP) assay [4] the starting state is a previously bound receptor-ligand complex (either in states *C*_0_ or *C*_1_) that, due to an externally applied tensile force, once the bond is broken, even if the two molecules keep the proper orientation, the probability of rebinding is very low. This experimental setup can be captured more accurately with the following diagram:

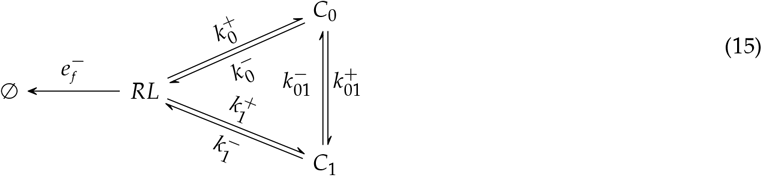

where 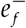 and the other rates are constant rates modulated by the applied force.

### 3.3. Conditions for the existence of catch-slip bonds in BCR-Ag interactions

In cell-to-cell experiments using the BFP assay [4], one has 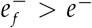 and *e*^+^ = 0 for *f* > 0. In simple terms, this means that by means of a mechanical procedure once a BCR unbinds from its ligand the two cell membranes immediately retract from each other. In this case, the interesting observable is the so-called bond survival distribution, *S*(*t*), that gives the distribution of empirically observed bond lifetimes at time *t* under conditions that prevent rebinding. Two time-scales in the distribution *S*(*t*) have been experimentally observed so that such bond survival distribution could be fitted phenomenologically to a sum of two exponentials, with weights 0 ≤ *ω*_1_ ≤ 1 and *ω*_2_ = 1 – *ω*_1_ and respective rates *k*_1_ and *k*_2_ [55]. As shown in Appendix A1 this empirical result can be formally derived assuming that the bond survival distribution is a quasi-stationary distribution [56]. Thus, the survival distribution is given by:

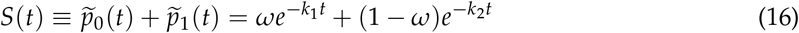

with

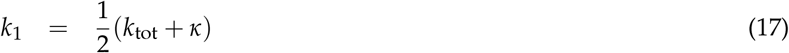

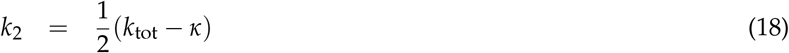

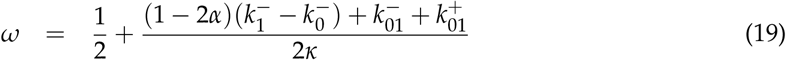

where

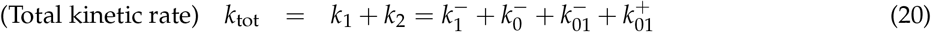

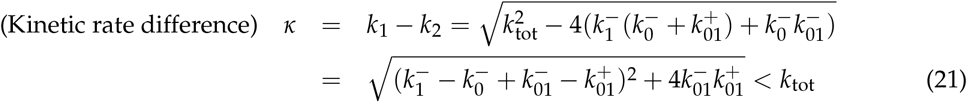

and *α* and (1-*α*) are the initial occupancy probabilities, respectively, of states *C*_1_ and *C*_0_. From Eqs. (A4)–(A5) it turns out that *α* boils down to the simple expression:

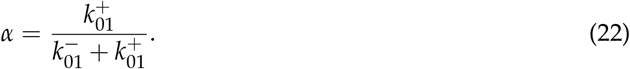

The importance of Eqs. (16)–(22) is that they allow us to explore the effect of the force on the bond survival distribution *S*(*t*). Specifically, if the bond kinetics is understood under an allosteric model where the kinetic rates depend on the free-energy landscape containing the work done by the applied force *f*, then, in our notation, one has [48]:

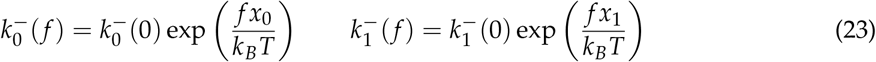

and

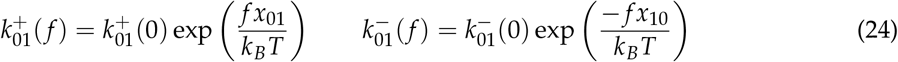

where *k_B_* is the Boltzmann constant, *T* is the absolute temperature of the system, and *x*_0_ and are the distances from the minima of the receptor-ligand complex potential energy, respectively, in the *C*_0_ and the *C*_1_ states to the maximum of the potential energy that acts as a barrier between the bound and the unbound (*RL*) states. Similarly, *x*_01_ and *x*_10_ in Eq. (24) represent the distances or widths from the minima in the *C*_0_ and *C*_1_ states, respectively, to the maximum of the potential energy between both states.

Equations (23)–(24) summarize the changes produced by the force in the free energy landscape relating states *RL, C*_0_ and *C*_1_ in Eq. (15). Note that the effect of the force is different on each rate and this can explain why some systems display catch-bond behavior while others do not. Nevertheless, this outcome depends on how *k*_1_, *k*_2_ and *ω* change with the applied force. More concretely, note that the relative weight of each timescale in Eq. (16) (quantified by *ω*) would also depend on the applied force. This effect might be observed experimentally because the expected time *t_x_* at which the survival function transitions from one timescale to the other depends on *κ* and *ω* and, hence, indirectly on the applied force *f*. This can be easily seen by considering that the time *t_x_* is determined by the condition *ωe*^−*k*_1_*t_x_*^ = (1 - *ω*)e^−*k*_2_*t_x_*^, which yields after some algebraic manipulation:

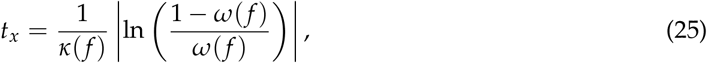

where *κ* = *k*_1_ – *k*_2_. Note that *t_x_* does only exist for force values at which *ω*(*f*) ≥ 1/2. Interestingly, some recent studies suggest that at least at *f* = 0 the transition rates between *C*_0_ and *C*_1_ are faster than the unbinding rates into *RL* [54], that is,

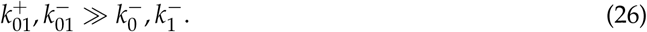

If all the distances in Eqs. (23)–(24) are within the same order of magnitude, namely, *x*_0_ ≃ *x*_1_ ≃ *x*_10_ ≃ *x*_01_ ≡ *x*, then for *f* > 0 the above relationships still hold and 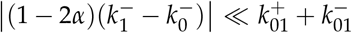, and hence Eq. (19) can be approximated by:

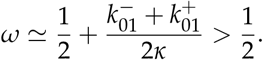

In general, the system’s response to the force will be determined by the sensitivity of the parameters with respect to the applied force. The sensitivity *σ_X_*(*f*), of the variable *X* with respect to *f* [57,58] is defined as:

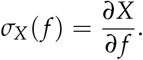

Applying this to the rates in Eqs. (23)–(24) we find that

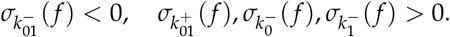

From Eq. 22, and taking into account that for *f* = 0 the state *C*_0_ is more favorable, it follows that *α*(0) < 1/2 and therefore 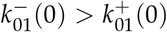. Hence, we can assume the following hierarchy of kinetic rates

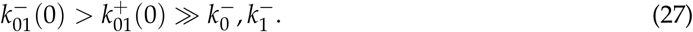

This means that, for small forces, the (negative) sensitivity of 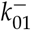 will dominate for lower values of the force as long as, roughly, (i) all the distances in Eqs. (23)–(24) (*x*_0_, *x*_1_, *x*_01_ and *x*_10_) are of the same order, and (ii) 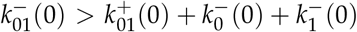 (what is plausible taking into account our previous discussion).

Under these reasonable conditions, we can conclude that,

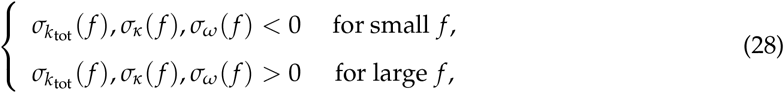

what provides a mechanism for the qualitative behavior of the survival function defined in Eq. (16). This situation is depicted schematically in Fig. 4.

**Figure 4.**
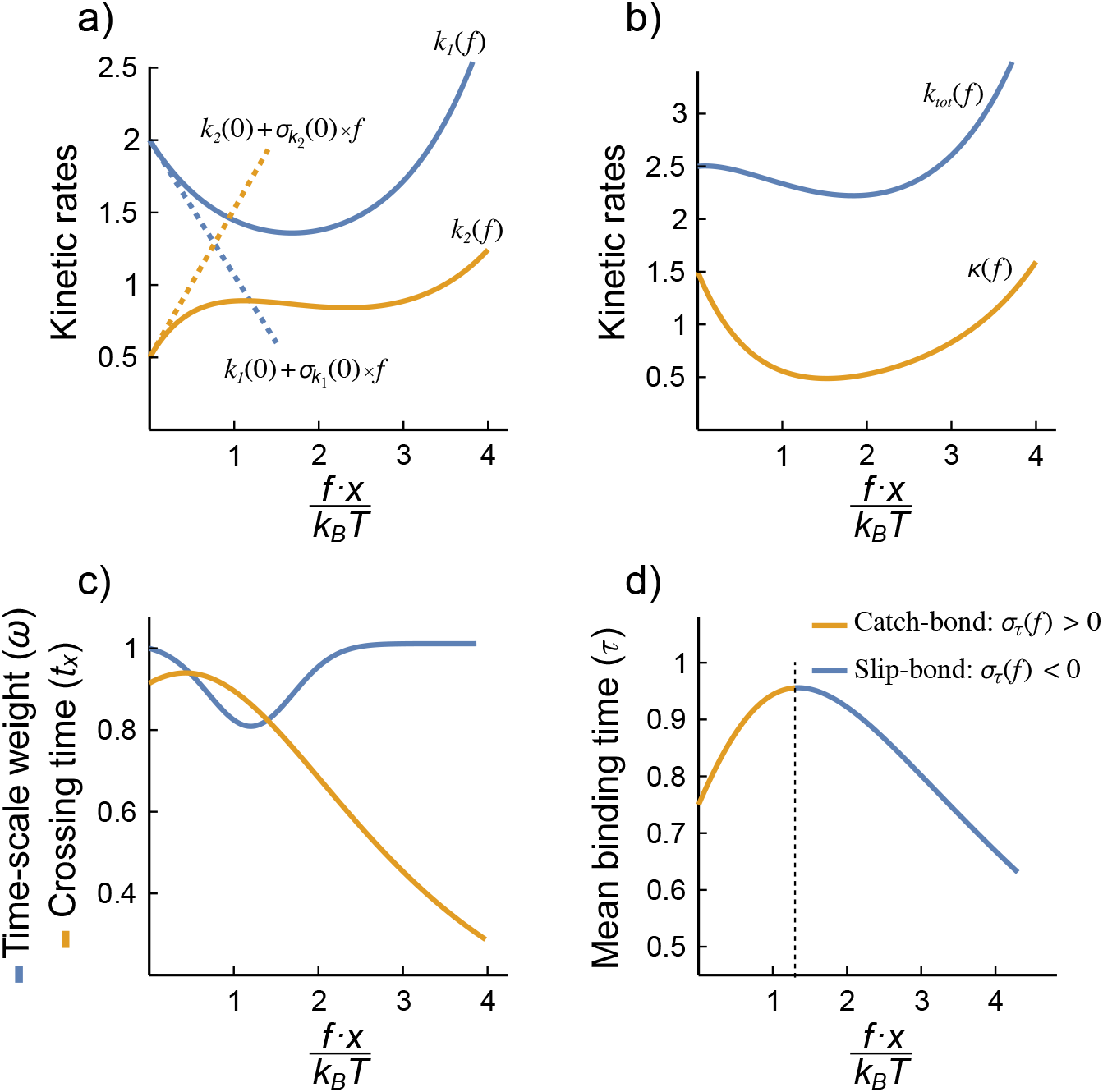
The BCR-Ag interaction might display a catch-bond behavior. Under the rather generic conditions outlined in the main text, the dependence on *f* of quantities like the main kinetic rates, the rates *k*_tot_ and *κ*, timescales weight *ω*, and times *t_x_* and *τ* can be captured. a) Variation of the main kinetic rates, *k*_1_ and *k*_2_, with the (scaled) applied force; dashed lines are the corresponding tangent lines at *f* = 0 with slopes equal to the rates sensitivities at *f* = 0. Note how the negative sensitivity of 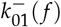 affects the sign of the sensitivity of *k*_1_ and *k*_2_ at low forces. b) Total kinetic rate, *k*_tot_ and kinetic rates difference, *κ*. c) Weight of the timescales, *ω*, and crossing time, *t_x_*. d) Dependence of the mean bond lifetime, *τ*, on the scaled applied force. The combination of *k*_1_, *k*_2_ and *ω* in Eq. (29) and the negative sensitivity of 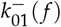 opens the possibility of a catch-bond behavior (the binding time increases with the applied force, at least for low forces). These plots are qualitative and the units are arbitrary. In practice, the shape of the curves depend on the specific values of the kinetic rates at zero force and the distances, *x_k_*, in Eqs. (23)–(24).

This change in the qualitative behavior of the survival function with force has been observed in platelet adhesion [55], and therefore it is sensible to experimentally test it in BCR-Ag interactions. If observed, it would support the idea that the BCR-Ag interactions are subject to force-dependent kinetic rates and also would help to estimate *x*01 and *x*10 in Eq. (24).

Equation (16) also allows us to explore the dependence of the mean bond duration and the applied force. Note that the probability density of the bound state as a whole (*C*_0_ + *C*_1_) is given by,

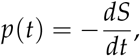

so the mean bond duration is simply

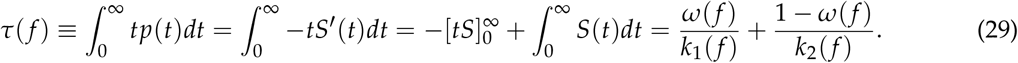

From this expression, it is straightforward to show that

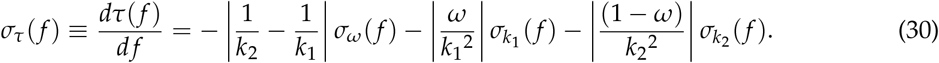

Hence, the sensitivity of the mean time is the sum of negative *weights* times the sensitivities of *ω*, *k*_1_ and *k*_2_.

The above assumption on the similarity of the order of magnitude of all the distances in Eqs. (23)–(24) implies that the scaled force, *fx*/*k_B_T*, can take values below a positive threshold,

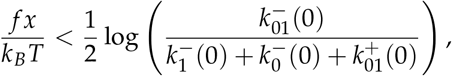

such that the timescales *k*_1_ and *k*_2_, their respective weights *ω* and (1 - *ω*), and the mean binding time *τ* would display non-monotonic behavior as a function of the scaled force, as shown in Fig. 4. In particular, note how *τ*(*f*) would increase for small values of the (scaled) force —catch bond behavior— and later it would decrease —slip-bond behavior.

This deserves a more thorough (qualitative and quantitative) exploration and it is out of the scope of this Perspective article. However, it is worth emphasizing that a role for catch-bonds in BCR-ligand interactions is clearly an open and not unlikely possibility.

## 4. Discussion

We have argued here that Ab-Ag binding constant rates measurements, both 3D and 2D, are effective estimations that may be not even proportional to the intrinsic binding rates. We have also shown that effective estimations of 3D constant rates may be not proportional to the effective 2D constant rates and, hence, they can be misleading when used to interpret results involving 2D BCR-Ag interactions, particularly BCR-driven B-cell activation and/or Ag endocytosis.

Moreover, irrespective of the effect different isotype glycosylations may have on the constant rates of Abs and BCRs [29], we have clarified here that Ig class switching and Ag in solution might significantly alter the effective binding constant rates of BCR-Ag and Ab-Ag interactions, mainly because of the concomitant change in the rotational diffusion rate constants. This is particularly relevant to interpret correctly —in terms of affinity maturation— the 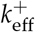 and 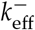 of Abs and BCRs of GC B cells that have undergone class switch. In particular, using 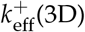 and 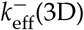 as surrogates, respectively, of *k*^+^ and *k*^−^, and according to what was discussed in subsection 2.4 with respect to 3D Ab-Ag interactions, there are two main, opposite possibilities: 1) Class switch of a BCR brings an increase of the 3D *k*^+^-threshold 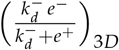. This increase will cause in the corresponding secreted Abs a decrease of the effective constant rate 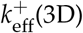 but an increase of 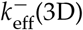 (see Appendix A.2), in spite of having the same variable region than before it switched. If such class-switched BCR mutates such that it increases *k*^+^ or decreases *k*^−^, this could be detected by SPR as an increase of 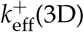 or a decrease of 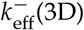, respectively. In other words, a BCR class switch that causes an increase of the 3D *k*^+^-threshold may allow detecting mutation-driven maturation of secreted Abs. 2) Class switch causes a decrease of the 3D *k*^+^-threshold, with a corresponding increase of 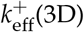 and decrease of 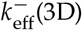. In this case, since 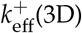 will get closer to the 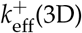 plateau, a mutation-driven increase of *k*^+^ could be barely detected or not at all by the SPR technique in the corresponding secreted Abs, and the concomitant decrease of 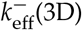 could not correspond to a decrease of *k*^−^, but to a decrease of the affinity dissociation constant *K_D_*.

In contrast, as we have discussed above (subsection 2.4), Ag barely contribute to translational and rotational diffusion in 2D BCR-Ag interactions, and only class switch of BCRs from IgD to IgG subclasses impacts significantly the translational diffusion constants (see subsection 2.2), because the corresponding IgD and IgG values differ nearly 10-fold, and the rotational diffusion constants. Thus, a BCR class switch from IgD to IgG1 can increase the *k*^+^-threshold 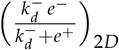 by 8–10-fold. However, other BCR class switches will affect only the BCR rotational diffusion constant rates, and we expect the corresponding change in *k*^+^-threshold to be small as shown in the following. Let us take for 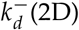 the upper bound 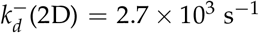, and consider a BCR with 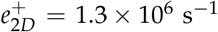 and 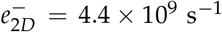. A BCR class switch from IgG1 to IgG2b can lead to a 2-fold increase in *D_t_*, a 1.5-fold increase in *D_a_* and a 1.5 increase in the tilt angle *θ*, which implies a 3-fold increase in *e*^+^(2D) and a 1.6-fold increase in *e*^−^(2D). A simple calculation shows then that the *k*^+^-threshold would decrease from 8.8 × 10^6^ s^−1^ to 4.8 × 10^6^ s^−1^, which is less than a 2-fold change. Therefore, whether *k*^+^ ≪ 2D *k*^+^-threshold or *k*^+^ 2D *k*^+^-threshold before a BCR switch, that condition will likely persist after switching. This predicts that estimations of 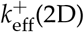 and 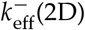 will not depend on the BCR isotype, except for a switch from BCR IgD.

When GC B cells bind an Ag presented by FDCs they form an immune synapse that later retracts, pulling the BCR-bound Ag. This Ag pulling, when succesful, ends up creating an endosomal vesicle containing the extracted material [43,59–61]. If the multivalent binding of immune complexes to Fc and complement receptors on FDCs is stronger than the rupture force of FDC membrane integrity nearby those immune complexes, then it can be expected that, in general, the tensile force either breaks the BCR-Ag bond or it extracts bites of the FDC membrane, including immune complexes plus Fc and complement receptors. Otherwise, the tensile force will lead either to a break of the BCR-Ag bond or to extract only immune complexes. In principle, the force generated by this Ag pulling can modify the BCR-Ag bond strength either decreasing it (*slip*-bond), or first increasing it and then decreasing it for increasing forces (*catch-slip* bond) [4,62]. Some recent observations lend support to this second possibility [43,50,63]. In this later case, the increased force resistance can reach the point at which a large Ag extraction takes place. Moreover, increasing the lifetime of a bond when a tensile force is applied could facilitate extracting bites of the antigen presenting cell membrane together with Ag complex. Indeed, this has been recently reported for B cells [43], reminiscent of an older report that showed how T cells can extract and internalize cognate pMHC molecules from antigen presenting cells [64].

During a moderate Ag extraction most of the engulfed material would be BCR-Ag complexes. Once processed in the endocytic vesicles, they will generate a large fraction of Ag-derived peptides, which will be presented to follicular T helper (Tfh) cells in form of peptide-MHC class II (MHCII) molecular complexes. In contrast, a strong Ag extraction would likely include portions of the FDC membrane [43,47], recruiting irrelevant proteins together with BCR-Ag complexes into the same endocytic vesicle. During the ensuing proteolysis in the endocytic vesicles a plethora of different peptides will be generated, many of them derived from FDC-membrane self-proteins, effectively diluting Ag-derived peptides in the vesicle. Consequently, the number of MHCII molecules presenting Ag-derived peptides would decrease. This implies that a B cell with high-affinity BCRs that bind Ag with slip bonds can present as many Ag-derived peptides as a B cell with moderate-affinity BCRs that bind Ag with catch bonds. Conversely, B cells with high-affinity BCRs that bind Ag with catch bonds can present as many Ag-derived peptides or even less than B cells with moderate-affinity BCRs that bind Ag with slip bonds, thus setting an effective trade-off to the strength of BCR-Ag bonds.

In summary, given the evidence supporting the existence of Ig allosteric states [16,52–54], we have shown here that under very broad conditions for the kinetic rates there is ample room for the existence of catch bonds in BCR-Ag interactions. If that is the case, then it could be expected that there would be some optimum BCR-Ag affinity for best Ag-derived p-MHCII presentation to Tfh cells. Moreover, if different BCRs with specificity for the same Ag differ in their bond properties (slip *vs* catch), then this property could be a selectable trait in GCs, favoring both high-affinity BCRs with slip bonding and moderate-affinity BCRs with catch bonding. So far, the immunological relevance of this possibility is unclear but the technology to confirm its existence is already available.

## Author Contributions

All authors contributed equally to this work.

## Acknowledgments

The authors thank Dr. Emilio Faro for his help in clarifying some equations.

## Abbreviations

The following abbreviations are used in this manuscript:

BCR: B-cell receptor
TCR: T-cell receptor
FDC: Follicular dendritic cell
GC: Germinal center
SPR: Surface Plasmon Resonance
BFP: Biomembrane Force Probe

## Appendix A Some mathematical results

### Appendix A.1 Master equations

The stochastic dynamics of Eq. (15) is captured by the solution of the system of differential master equations:

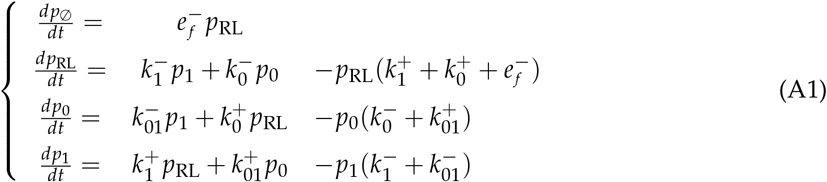

with 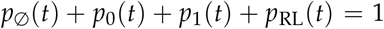. Note that the system of Eqs. (A1) includes an irreversible state (so that 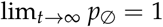).

The bond survival distribution can be computed as the probability of being in any state *C*_0_ or *C*_1_ *conditioned* to eventual decay into RL, respectively, 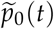 and 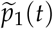:

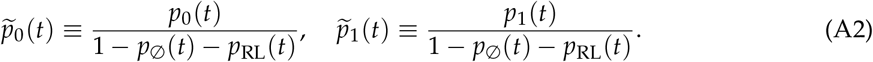

Thus,

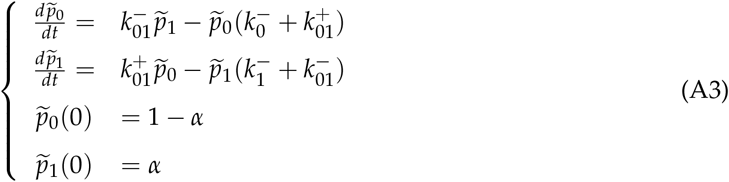

The solution to A3 is summarized in Eq. (16).

The initial condition *α* is the relative occupancy of state *C*_1_ at *t* = 0, so that from Eqs. (15) and (A1) and assuming *C*_0_ and *C*_1_ are in quasi-steady state,

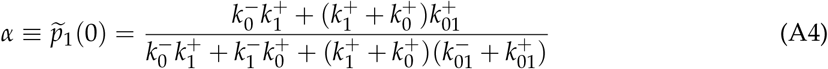

Assuming also that the reactions in Eq. (15) obey *detailed-balance* [23,65], namely,

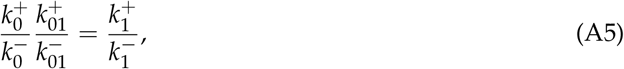

then we arrive at Eq. (22).

### Appendix A.2 Effect of k^+^-threshold increase (or decrease) on 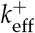 and 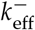

In general, given a ratio 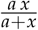 and a parameter *b*, with *a, x, b* > 0, one has:

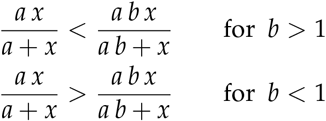

Given the equations:

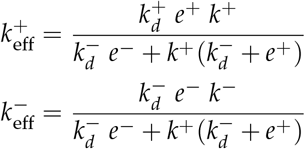

and considering that 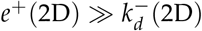 they can be re-written as:

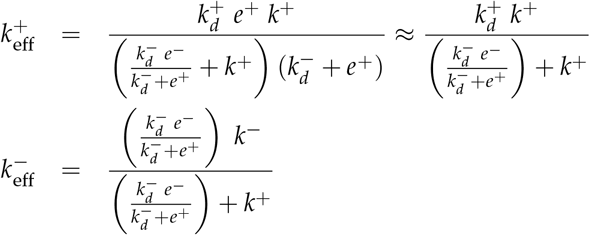

therefore, if 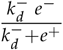 increases 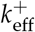 decreases and 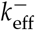 increases, and vice versa.

